# Efficient CRISPR/Cas9 mediated large insertions using long single-stranded oligonucleotide donors in *C. elegans*

**DOI:** 10.1101/2022.09.26.509570

**Authors:** Matthew Eroglu, Bin Yu, Brent Derry

## Abstract

Highly efficient generation of deletions, substitutions, and small insertions (up to ∼150 bp) into the *C. elegans* genome by CRISPR/Cas9 has been facilitated by use of single-stranded oligonucleotide donors as repair templates. However, efficient insertion of larger sequences such as fluorescent markers and other functional proteins remains inefficient due to lack of standardized methods for generating repair templates and labor intensive or cost prohibitive synthesis. We have optimized the simple and efficient generation of long single-stranded DNA for use as donors in CRISPR/Cas9 using a standard PCR followed by an enzymatic digest by lambda exonuclease. Comparison of long single-stranded DNA donors to previously described methods using double-stranded DNA yields orders of magnitude increased efficiency for single-stranded DNA donors. This efficiency can be leveraged to simultaneously generate multiple large insertions as well as successful edits without use of selection or co-conversion (coCRISPR) markers when necessary. Our approach complements the CRISPR/Cas9 toolkit for *C. elegans* to enable highly efficient insertion of longer sequences with a simple, standardized and labor-minimal protocol.

## Introduction

Currently, the most efficient method of genome editing in *C. elegans* is by injection of Cas9 protein-guide RNA (RNP) complexes into the syncytial germline (Cho et al., 2013; Paix et al., 2015). Precise and efficient edits such as deletions and small insertions can be obtained by provision of single-stranded oligonucleotide donors (ssODNs) which are commercially synthesized. Studies from worms to mammals have demonstrated the advantages of using single-stranded (ssDNA) donors over double-stranded DNA (dsDNA) in both efficiency and cytotoxicity (Levi et al., 2020; Wang et al., 2013; Zhao et al., 2014). However, the per base cost of outsourcing ssDNA synthesis remains prohibitive beyond ∼200 nucleotides for most applications.

Commonly used strategies for larger insertions employ linear dsDNA or plasmid donors, which often yield very low (<1%) insertion frequencies depending on locus and are toxic when injected at high concentrations. Alternatively, melting and annealing dsDNA donors (Ghanta and Mello, 2020) as well as addition of 5’ modifications (Ghanta et al., 2021) have been reported to increase insertion efficiencies. However, these methods involve specific annealing conditions difficult to standardize, laborious gel purification of donors, and added costs associated with 5’ modifications. Furthermore, the mechanism by which these treatments increase efficiencies remains unclear, making their optimization difficult.

Several methods have previously been described for the *in vitro* generation of long ssDNA fragments. The simplest method is a one-step asymmetric PCR which involves use of unequal primer concentrations during PCR to yield an excess of one product (Gyllensten and Erlich, 1988). Alternatively, a two-step method involving first a regular PCR reaction with one 5’ phosphorylated primer followed by digestion of the phosphorylated strand by lambda exonuclease, a strand specific nuclease, has also been reported to yield large amounts of ssDNA (Kujau and Wölfl, 1997). Both methods have been used to generate donors for use in CRISPR/Cas9 (Inoue et al., 2021; Kanca et al., 2019), although each presents challenges such as isolation of enough intended product at high enough concentrations and purities for use as donors. Furthermore, for longer insertions, systematic comparisons of ssODNs to double-stranded repair templates remains lacking.

Given the advantages of using ssODNs for generating smaller edits by CRISPR/Cas9 in *C. elegans*, we explored the use of such donors for larger edits and standardized their use for this application. Comparison of the simple methods used for generating long ssDNA revealed that the lambda exonuclease protocol consistently and rapidly (within <3 hours) yields high amounts (micrograms) of long ssDNA appropriate for use as donors in CRISPR/Cas9 mediated insertions. When compared to dsDNA and the recently described protocol using melted dsDNA, long ssODNs yield orders of magnitude greater efficiencies for larger inserts across a variety of loci with templates including fluorescent tags as well as enzymatic fusions. Overall, up to 92% of marker positive worms can display successful insertions depending on locus and injection, bringing the efficiency of long insertions to levels comparable to smaller edits. We leverage this efficiency to simultaneously knock in multiple large inserts in one round of injections.

## Results

### Generation of single-stranded DNA

To determine the simplest and highest yield protocol for generation of long ssDNA, we compared asymmetric PCR to lambda nuclease digestion (Fig. 1A). Repair templates encoding three different inserts (mScarlet, GFP and TurboID) at various genomic loci (*gld-1, rec-8* and *ccm-*3) were designed with up to 140bp homology arms on both ends of the insert and synthesized commercially as dsDNA at the lowest scale. Asymmetric PCR was performed on this template by varying the ratio of forward to reverse primers. In contrast, the template for the nuclease method was amplified with a PCR reaction using one 5’ phosphorylated primer to generate 10 µg of dsDNA. After spin column purification, a digestion was performed with lambda exonuclease followed by a final spin column purification. Both protocols yielded appreciable single-stranded DNA which could be visualized by gel electrophoresis (Fig. 1B-F). While asymmetric PCR was simpler, involving a single step, it often yielded nonspecific products at unintended lengths or unpredictable yields of ssDNA requiring template specific optimization (Fig. 1E, F). In contrast, digestion by lambda exonuclease consistently yielded a product at the expected size regardless of template (Fig. 1B-D). We therefore optimized the conditions for digestion and recovery of ssDNA using the lambda exonuclease method.

**Figure 1:**
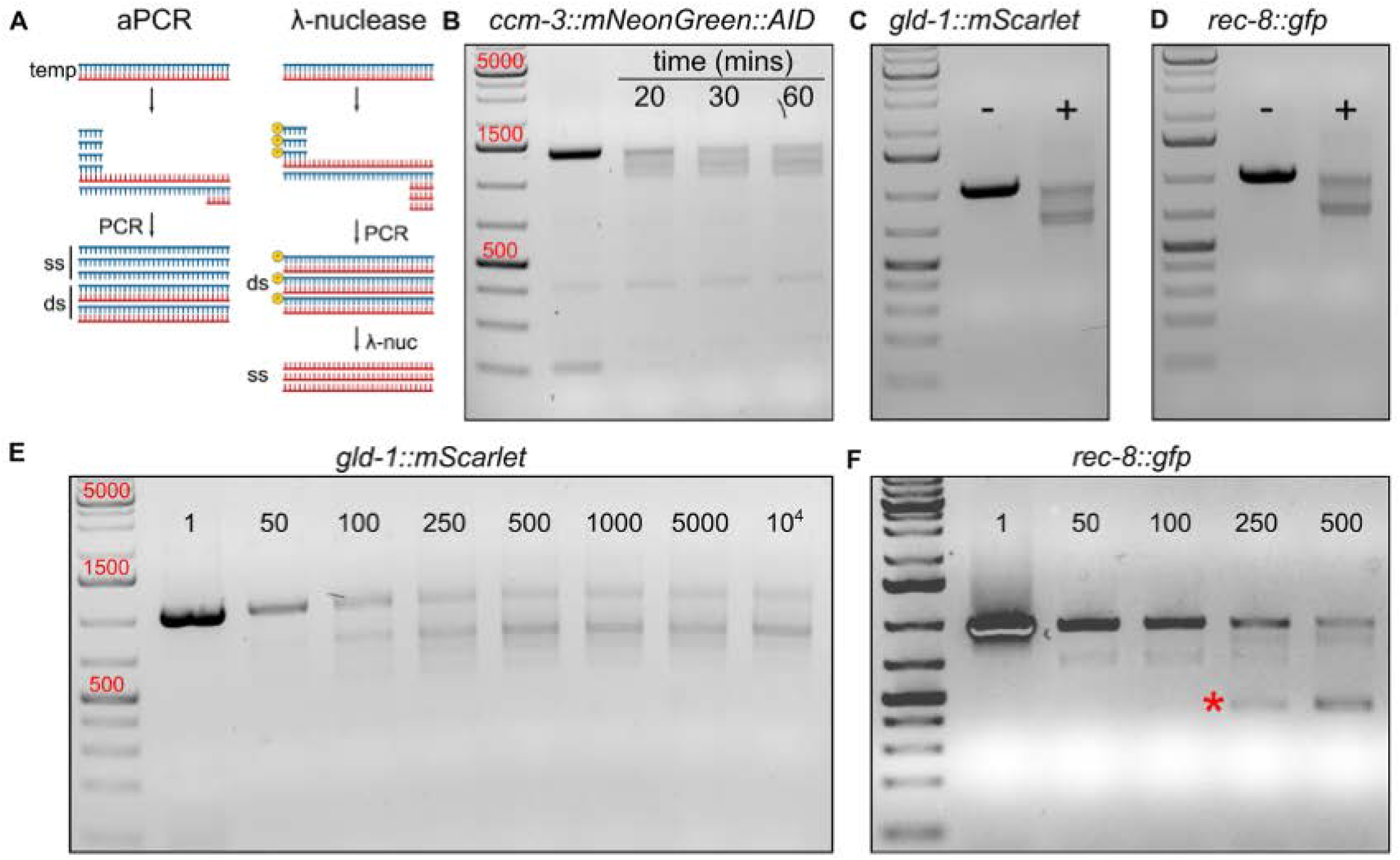
Comparison of asymmetric PCR and lambda exonuclease method for generating long ssODNs. **(A)** Schematic of asymmetric PCR (aPCR, left) and the lambda (λ) exonuclease method (right). In aPCR, different ratios of forward to reverse primer are used to generate an excess of one product. In the λ-exonuclease method, a normal PCR is first performed using one phosphorylated primer, followed by a subsequent digestion with λ-exonuclease which preferentially degrades the phosphorylated strand leaving ssDNA. **(B)** Time course of digestion with λ-exonuclease reveals that the reaction is complete after 30 mins incubation of 10 µg of a 1.4kb dsDNA fragment. **(C, D)** Similar results are obtained with two different templates, *gld-1::mScarlet* **(C)** and *rec-8::gfp* **(D)**. **(E)** Asymmetric PCR of the *gld-1::mScarlet* template reveals that ssDNA can be obtained using this method. However, unlike the λ-exonuclease method, multiple unintended bands are present. **(F)** Similarly, aPCR of *rec-8::gfp* yields significant amounts of an unintended product (star).

Time course of digestion showed that 10 µg of 1-1.4 kb dsDNA could be digested by 10 units of lambda nuclease within 30 minutes in a 50 µL reaction at 37°C (Fig. 1B-D). Rapid (5-10 min) purification of ssDNA was achieved with recovery yields >60% using spin columns (Monarch PCR & DNA Cleanup Kit (5 µg), NEB) with a modified protocol provided by the manufacturer described in Supplemental File 1.

### Long single-stranded donors yield highly efficient insertions

We compared the efficiency of insertions obtained using dsDNA, melted dsDNA as described previously (meltDNA), and ssDNA. Injection mixes were prepared as described (Paix et al., 2015) with minor modifications using a *dpy-10* coCRISPR marker along with a single guide RNA to the target locus and the long ssODN donor encoding the insert of interest (Fig. 2A). Successfully edited progeny were recovered as dumpy (*dpy-10* homozygous edits) or roller (*dpy-10* heterozygotes). Marker positive progeny were genotyped by PCR using one primer internal to the edit and one outside. Using dsDNA or melted dsDNA yielded similar editing efficiencies across 3 different loci (mean 7.5%, median 3.1%, n = 480 worms; and mean 3.0%, median 2.1%, n = 432 worms, respectively, Fig. 2B-D and S1-3). In contrast, ssDNA donors yielded significantly higher efficiencies (mean 39.9%, median 35.4%, n = 528 worms, Fig. 2B-D, S1-3). Furthermore, depending on locus up to 92% (*gld-1*) of marker positive progeny were edited representing a 100-fold increase in insertion efficiency compared to the alternative methods (Fig. 2B, S1), and at minimum efficiency was 12.5% while alternative protocols sometimes yielded no successfully edited marker positive worms (Fig 2B-D, S3). Among three ssDNA donor concentrations (500, 250 and 100 ng/µL), 250 ng/µL and above yielded comparable editing frequencies for *gld-1* (Fig. S4).

**Figure 2:**
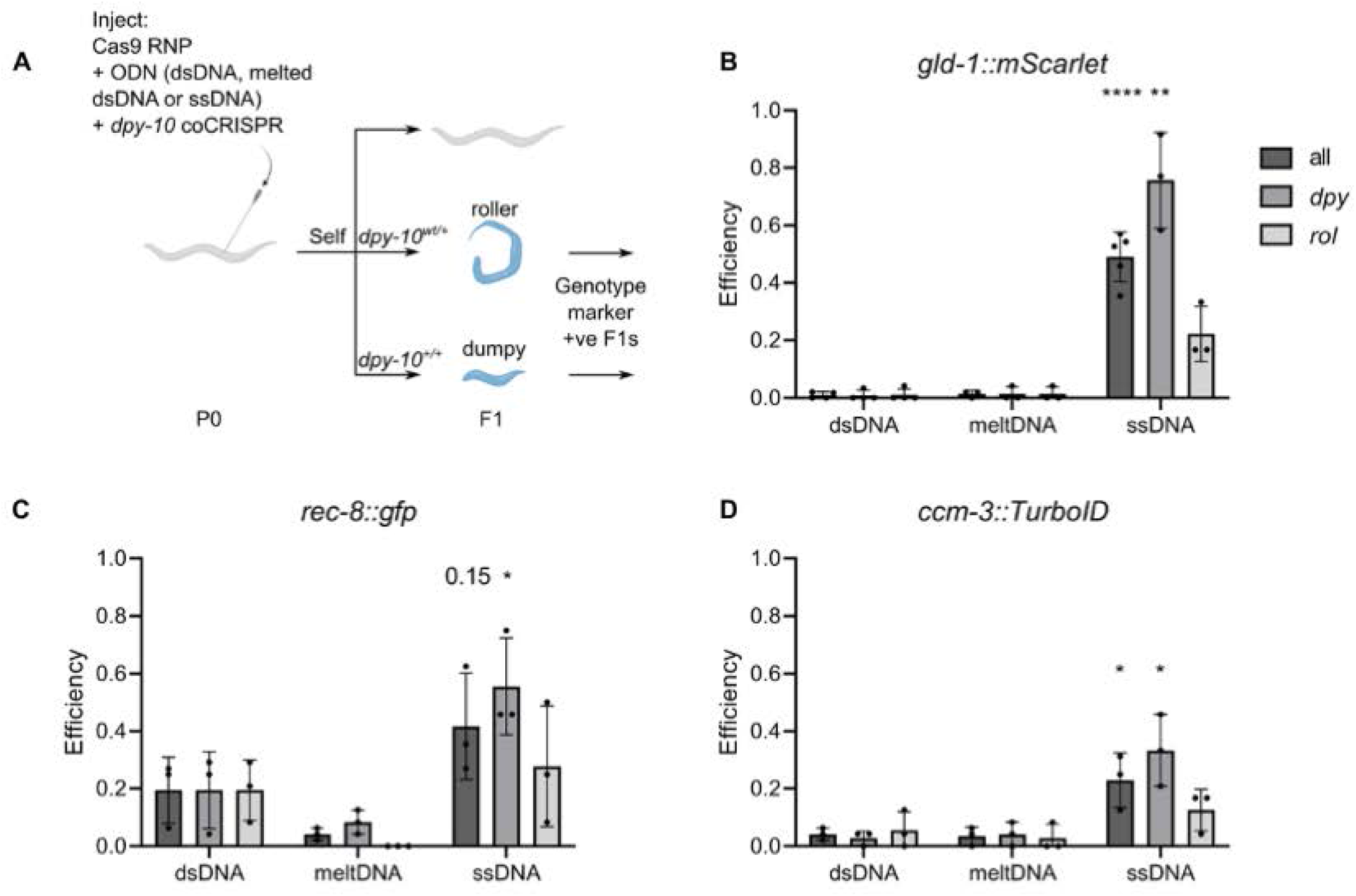
Long ssODNs enhance efficiency of edits across loci compared to alternative methods. **(A)** Schematic of injections used to score the efficiency of protocols. Gonads are injected with Cas9 ribonucleoprotein (RNP) complex, repair template and co-conversion marker (*dpy-10*). Roller and dumpy progeny are then screened by PCR for the intended insert. **(B)** Comparison of dsDNA, meltDNA, and ssDNA in generating a *gld-1::mScarlet* knock-in allele reveals significantly higher efficiency when using the ssDNA donor. Dumpy worms displayed the highest frequency of knock-ins when using ssDNA compared to rollers. Two-tailed t-test: **** *P <* 0.0001; ** *P <* 0.01. **(C)** Similar findings as **(B)** for *rec-8::gfp*. Two-tailed t-test: * *P <* 0.05. **(D)** Similar findings as **(B)** for *ccm-3::TurboID*. Two-tailed t-test: * *P <* 0.05.

The co-conversion marker *dpy-10* yields both biallelic edits in F1 progeny displaying a dumpy phenotype and monoallelic edits that develop as rollers. Rollers which are *dpy-10* heterozygotes are generally preferred as the *dpy-10* mutation can be selected out in a single generation, whereas *dpy-10* homozygotes must be outcrossed. We compared the efficiency of insertions in dumpy and roller progeny and determined that dumpy worms consistently showed the highest frequency of edits compared to rollers when using ssDNA donors (Fig. 2B-D). In contrast, dsDNA donors showed no correlation with the efficiency of the *dpy-10* edit. While isolating dumpy worms maximized insertion efficiencies, roller worms still displayed efficiencies greater than previous methods across loci (Fig. 2B-D).

### ssDNA donors enable double knock-in of large inserts

Given the efficiency of insertions generated by using ssDNA was high, we explored whether this could be leveraged to enable multiple knock-ins of large inserts. We coinjected the templates encoding *gld-1::mScarlet* and *rec-8::gfp* due to their ease of detection by fluorescence imaging and well characterized localizations, along with the *dpy-10* co-conversion marker (Fig. 3A). We genotyped *rec-8* first as it was the less efficient of the two based on prior injections (Fig. 2B, C). Among roller worms, 9/48 were positive for the *rec-8::gfp* insertion (Fig. 3B) whereas among dumpy worms 17/48 were positive (Fig. 3C). From the roller *rec-8::gfp* positive worms, 5 were also positive for some *gld-1::mScarlet* band (Fig. 3D), and among dumpy worms 7 were positive for *gld-1::mScarlet* (Fig. 3E). Although *rec-8::gfp* positive worms mostly displayed a single band at the expected size (Fig. 3B, C), *gld-1::mScarlet* genotyping revealed a variety of bands indicating partial or erroneous insertions (Fig. 3 D, E). This was in contrast to *gld-1::mScarlet* single insertions which mostly displayed one PCR product at the intended size (Fig. S1). We therefore assessed what percent of the double positive worms inserted both fluors correctly. Among double positive dumpy worms showing the expected PCR product for both *rec-8::gfp* and *gld-1::mScarlet*, 1/4 had inserted both correctly and displayed the appropriate fluorescence expression pattern (Fig. 3F). Among double positive rollers, all displayed a band at the incorrect size for *gld-1::mScarlet* indicating partial insertions. Overall, the efficiency of insertions was sufficient to simultaneously insert multiple long sequences within one generation, in addition to the point mutation co-conversion marker.

**Figure 3:**
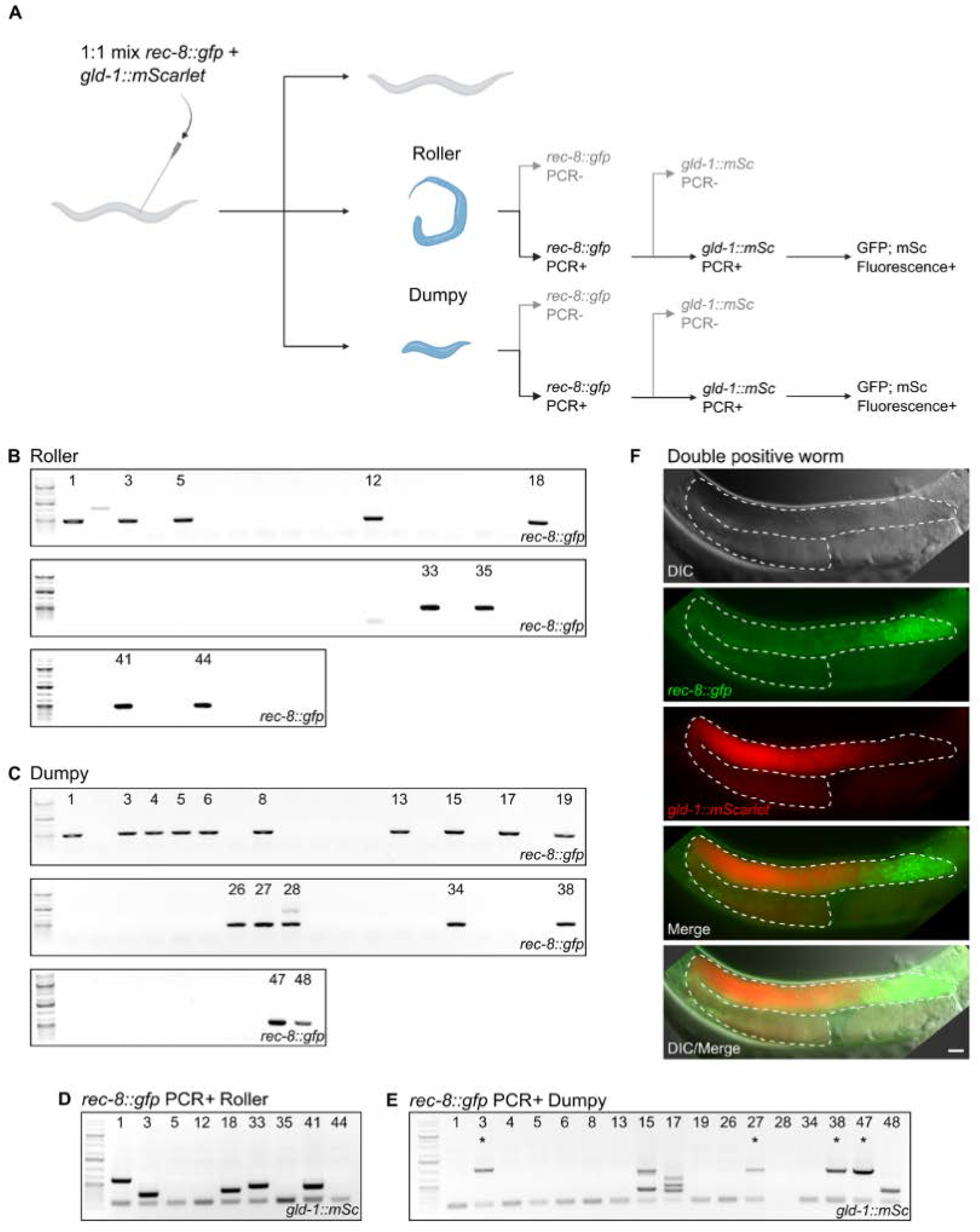
Use of long ssODNs to generate double knock-in. **(A)** Schematic of injection used to generate double knock-in of *gld-1::mScarlet* and *rec-8::gfp*. After injection, 48 roller and dumpy progeny were isolated and genotyped by PCR for *rec-8::gfp*. Worms which were positive for *rec-8::gfp* were subsequently screened for *gld-1::mScarlet*. Worms double-PCR-positive for the insertions were validated by fluorescence microscopy. **(B)** Roller worms screened by PCR for *rec-8::gfp*. Out of 48 rollers, 9 were successfully edited. **(C)** Dumpy worms screened by PCR for *rec-8::gfp*. Out of 48 dumpy worms, 17 were successfully edited. **(D)** Roller worms positive for *rec-8::gfp* were genotyped for *gld-1::mScarlet*. **(E)** Dumpy worms positive for *rec-8::gfp* were genotyped for *gld-1::mScarlet*. 4 worms displayed a *gld-1::mScarlet* product at the correct size (stars). **(F)** Representative double-edited worm displaying expression of *gld-1* and *rec-8* at the expected compartments of the germline.

## Discussion

Despite the widespread use of CRISPR/Cas9 for genome editing and the numerous protocols described in the literature, efficiently generating longer insertions such as fluorescent tags remains challenging. This issue may arise from the complexity of existing protocols describing steps hard to standardize like DNA annealing as well as laborious methods such as gel purification or addition of costly covalent modifications to donors. Indeed, we could not reproduce previously reported insertion efficiencies simply by melting dsDNA donors. To address these deficiencies, we generated a protocol that simplifies every step down to highly standardized and minimally laborious methods while increasing insertion efficiencies.

For small edits (indels) as well as any deletions, use of commercially synthesized ssDNA donors has been reported to be highly efficient, and labs routinely use such donors. However, chemical synthesis of longer ssDNA fragments remains cost prohibitive. Our comparisons of the different enzymatic methods for generating long ssDNA fragments revealed that both asymmetric PCR and lambda nuclease digestion can, in principle, generate significant amounts of ssDNA. While asymmetric PCR was more rapid, taking a single step, it also presents with several drawbacks that precluded its use by us. Firstly, there was no single forward to reverse primer ratio that worked well with all templates. More importantly, some templates yielded unexpected off-target products and there was no optimal primer ratio that alleviated this without sacrificing yield, necessitating labor-intensive gel purification for isolation of the intended product. In contrast, use of lambda exonuclease both resulted in high yields and predictable products, and this was consistent across all templates we used. Therefore, we recommend the lambda exonuclease method to generate ssDNA fragments for use when inserting longer fragments by CRISPR/Cas9-mediated genome editing.

When compared with alternative methods utilizing linear dsDNA PCR products, as well as melted dsDNA donors, we noted that ssDNA donors consistently yielded orders of magnitude higher editing frequencies across different loci. Previously established methods have pushed the efficiency of smaller edits by CRISPR/Cas9 to very high levels, frequently resulting in over half to nearly all progeny of injected parents displaying the desired edit. Certainly, for some loci such as *dpy-10* the high penetrance of visible phenotypes in progeny have enabled their use as selection markers of edited individuals. We have now achieved longer insertion frequencies at some loci comparable to those of smaller inserts (e.g., nearly all *dpy* worms for *gld-1*). Leveraging this efficiency, we have simultaneously generated dual knock-in of longer sequences at multiple independent loci.

Several considerations should be made when utilizing our method. While dumpy worms which are *dpy-10* homozygotes yield the highest insertion frequencies, it takes an additional outcrossing step to eliminate this marker. In contrast, while rollers displayed lower insertion frequencies compared to dumpy worms, it was still high enough to isolate numerous insertions in as little as 24 roller worms and still provided a significant advantage over previous methods. We routinely observe hundreds of rollers from a handful of (5-10) injected worms, and rarely find it necessary to screen dumpy worms for our insertions. Even with the double insertion, we isolated several rollers which displayed both PCR products, though none had inserted both functionally. In principle, scaling up the number of screened rollers should yield doubly edited worms with both constructs correctly inserted. In any case, when inserting multiple constructs simultaneously, additional care should be given as the frequency of partial or erroneous insertions appeared higher in our single proof-of-principle experiment. Nonetheless, the successful double insertion highlights the promise of this approach. Furthermore, in situations where *dpy-10* cannot be used as a marker, for instance when the desired insert is closely linked to *dpy-10* on Chromosome II, alternative markers may be necessary. We again leverage the efficiency of our method to describe a marker-free approach (Described in Supplemental File 1) enabling the isolation of successfully edited progeny without any co-conversion or extrachromosomal selection marker.

Overall, we offer a highly robust, standardized, efficient, cloning-free and labor-minimal protocol for obtaining large insertions by CRISPR/Cas9.

## Methods

### Nematode growth and maintenance

All nematode strains were maintained at 20°C on nematode growth medium (NGM) agar plates with OP50 *E. coli* as a food source (Brenner, 1974). All CRISPR/Cas9 mutant strains were generated in the Bristol N2 background.

### Microscopy

Worms were immobilized with 20mM tetramisole on 4% agarose pads and sealed with a coverslip. DIC and fluorescence microscopy was carried out using a Leica DMRA2 system equipped with DIC and fluorescence optics.

### Scoring insertions

Injected worms were plated on NGM and allowed to self-fertilize. Individual dumpy or roller self-progeny were lysed using 5uL worm lysis buffer (50mM KCl, 100mM Tris-HCl pH 8.2, 2.5mM MgCl_2_, 0.45% NP-40, 0.45% Tween 20). PCR was performed (1. 95°C for 5:00 mins; 2. 95°C for 20 secs; 3. 59°C for 15 secs, 4. 72°C for 1:00 min; 5. 34x cycle between steps 2-4; 6. 72°C for 2:00 mins; 7. Hold at 12°C) using 1uL of worm lysate and Advanced 2x HS-Red Taq PCR mastermix (Wisent Cat: 801-200-DM). One primer was designed to target a sequence internal to the insert whereas the other targeted outside the insertion. Agarose gel electrophoresis was performed in 1% gels and worms were scored as having insertions if bands were observed.

### CRISPR/Cas9

A detailed standard operating protocol is included as a supplemental file to this manuscript (Supplemental File 1). Cas9 protein was obtained from IDT (Alt-R® S.p. HiFi Cas9 Nuclease V3, 100 µg, Cat:1081060), aliquoted at 5mg/mL concentration and stored at -80°C until use.

## Data Availability

All additional raw data and reagents available on request.

## Contributions

M.E. conceived of the study and performed experiments. B.Y. performed CRISPR/Cas9 experiments. W.B.D. also conceived of the study and secured funding.

## Acknowledgements

We thank Dr. Oliver Hobert for his suggestion to submit this protocol for publication.

## Figure Captions

**Supplemental Fig. 1.**
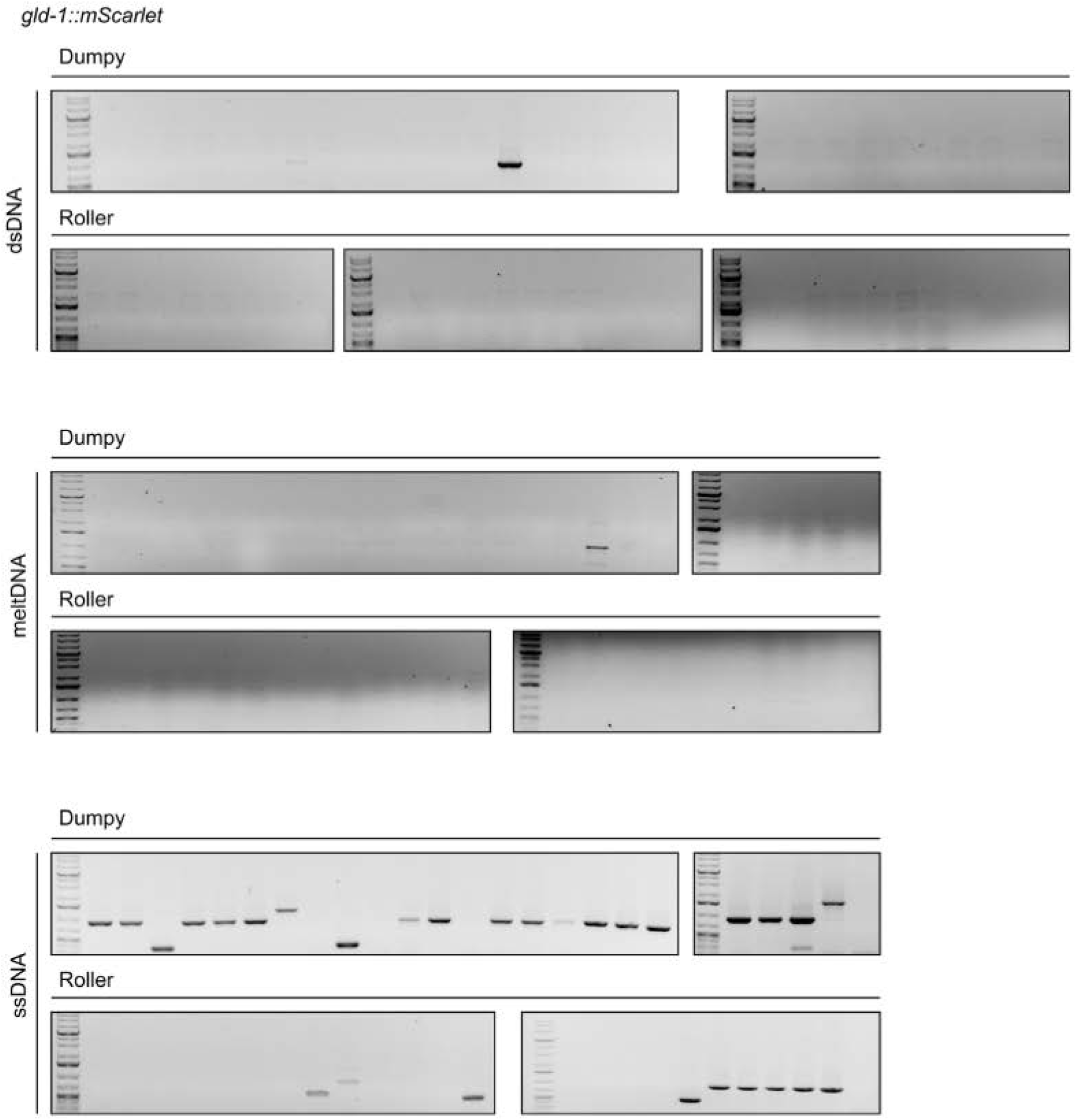
Representative gel images of *gld-1::mScarlet* using indicated repair template to generate the insertion.

**Supplemental Fig. 2.**
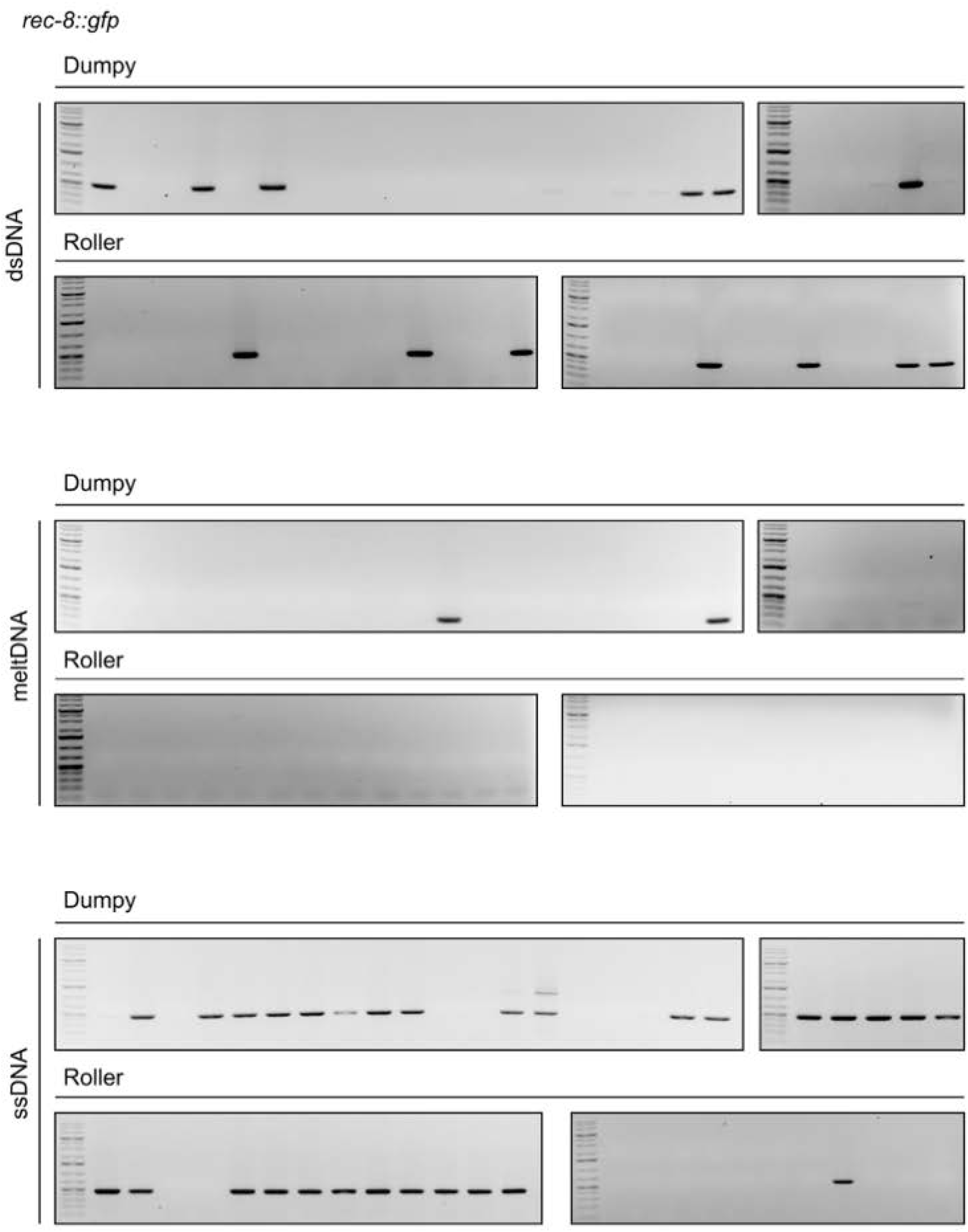
Representative gel images of *rec-8::gfp* using indicated repair template to generate the insertion.

**Supplemental Fig. 3.**
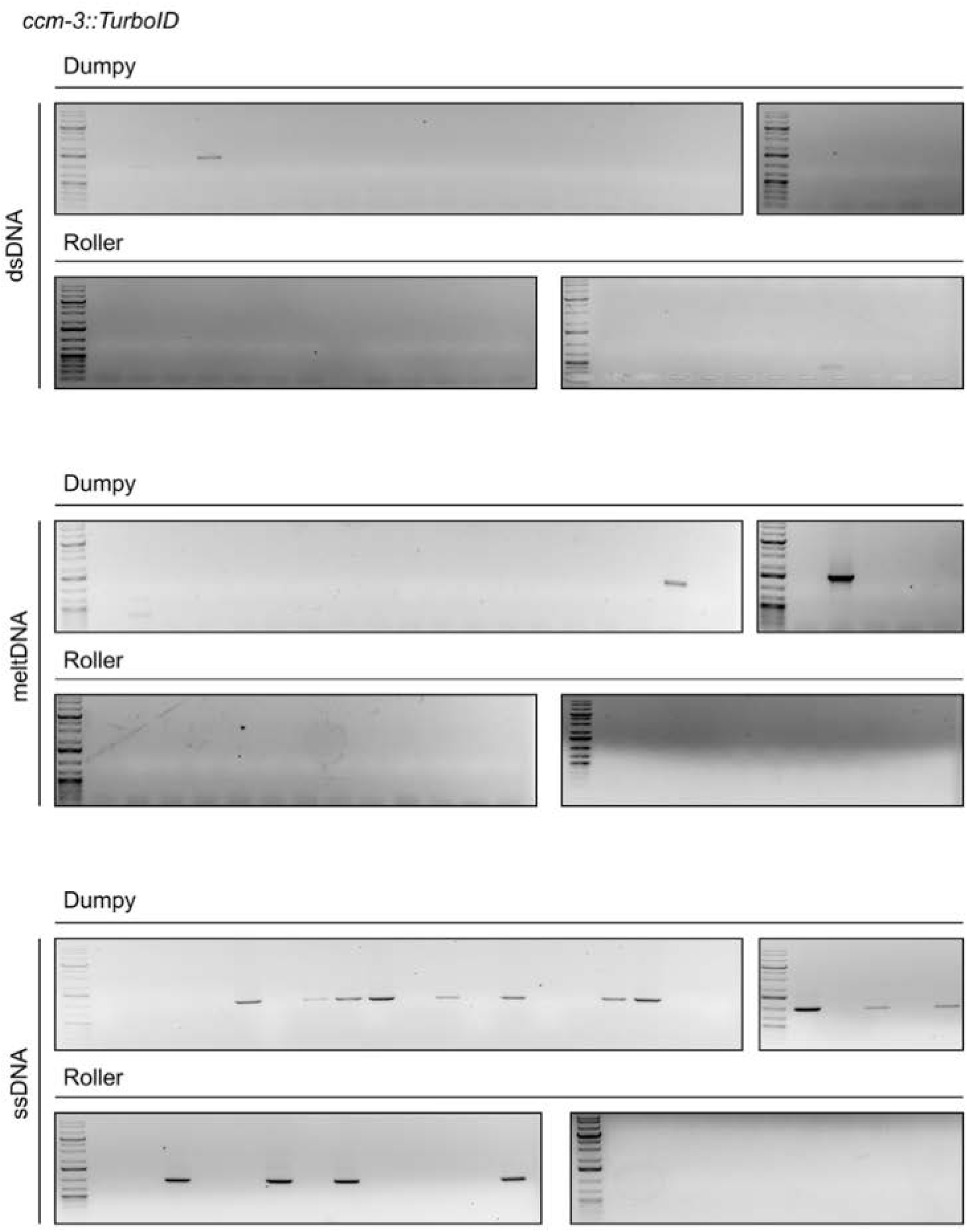
Representative gel images of *ccm-3::TurboID* using indicated repair template to generate the insertion.

**Supplemental Fig. 4.**
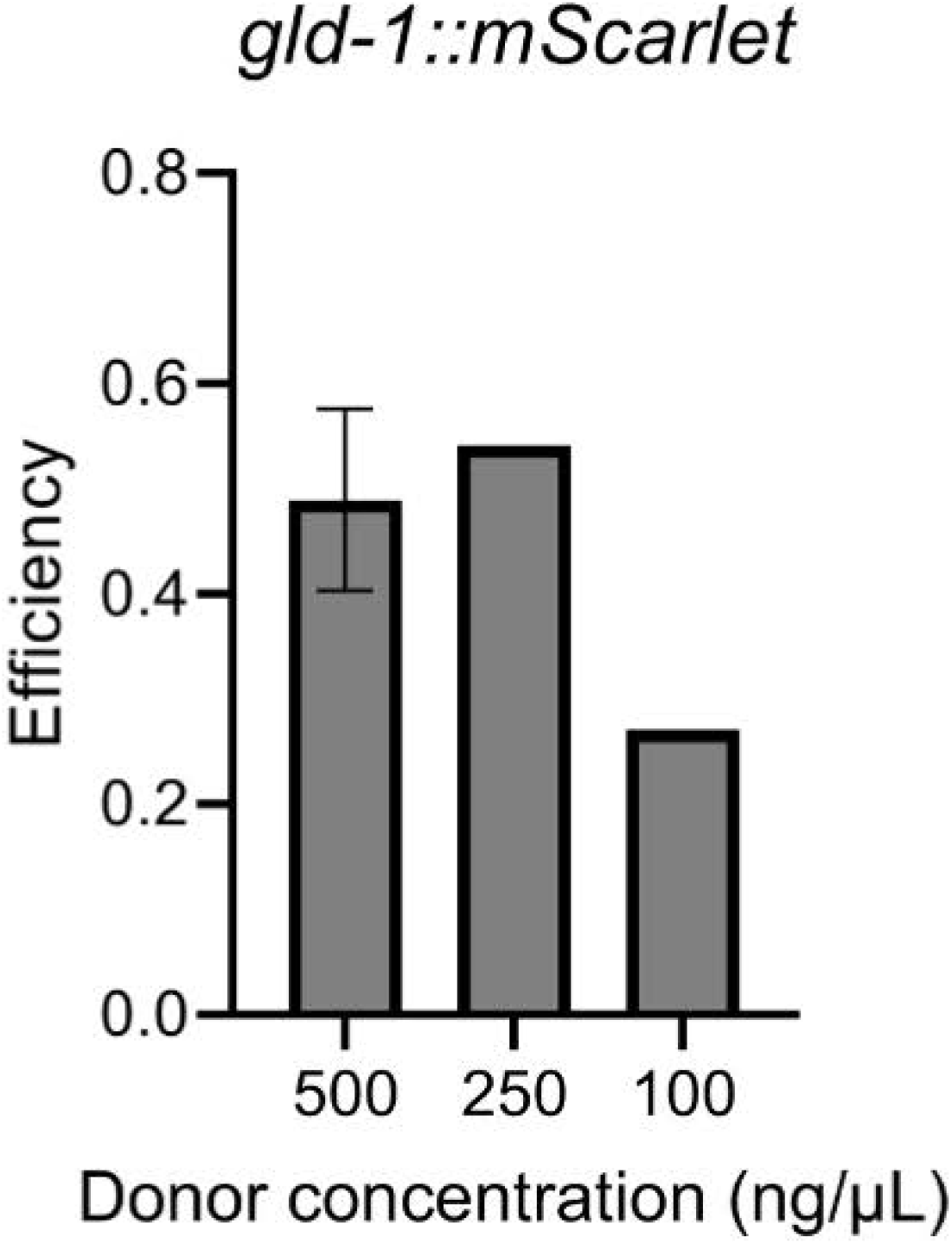
Titration of the ssODN donor concentration in the injection mix reveals similar editing efficiencies above 250 ng/µL.

## Genome editing by CRISPR/Cas9

A protocol adopted from the Seydoux lab is used (Paix *et al*, 2015, PMID: 26187122). In summary, guide RNAs, repair templates, and Cas9 RNP is injected directly into the germline. *dpy-10* is commonly used as a co-CRISPR marker.

### Deletions and small insertions

#### Designing and ordering guides

1. Design:
  - For deletion, two guides (crRNA) are used at each end of deletion.
  - For insertions, a single guide is sufficient.
  - Guides designed using IDT’s crRNA design tool
    - Query ∼250nt sequence that covers region of interest. Guides should target as close as possible to edit site.
2. Order guides from IDT (Alt-R CRISPR crRNA, 2nmol):

#### Designing and ordering repair template

1. Design:
  - For a deletion, design ssODN with homology arms on either side of desired deletion. Generally, 50-75nt on each side is sufficient.
  - For a small insert or SNP, e.g. 3xFLAG, add equal homology arms on each end of desired mutation sequence up to total 150nt.
  - Repair template should be on opposite strand as sgRNA.
  - If necessary, PAM sites should be mutated, or recognition sequence should be disrupted by insert to prevent repeated cutting by Cas9
2. Order ssODN (EXTREmer oligo from eurofins)

#### Preparation of crRNA and ssODN

1. Once all components have arrived:
  - Resuspend 2nmol crRNA in 20uL duplex buffer → 100 µM
  - Resuspend ssODN in indicated volume MG H_2_O → 100 µM
2. Mix all RNAs together
  - 0.68uL 200µM tracrRNA
  - 0.40uL 100µM *dpy-10* crRNA
  - 0.96uL each of two gene-specific guides, or 1.92uL one guide
3. Anneal in thermocycler (program is pre-set)
  - 95°C for 5 min
  - 10°C for 5 min
  - 4°C hold (or store at -20_o_C until needed)

#### Injection mix recipe

- 1.02uL annealed RNA mix
- Add 1.25uL Cas9 and incubate 5 min at room temp
- 0.20uL gene-specific ssODN (100µM)
- 0.67uL *dpy-10* ssODN (10µM)
- 1.36uL MG H_2_O (use more if less Cas9 is used)

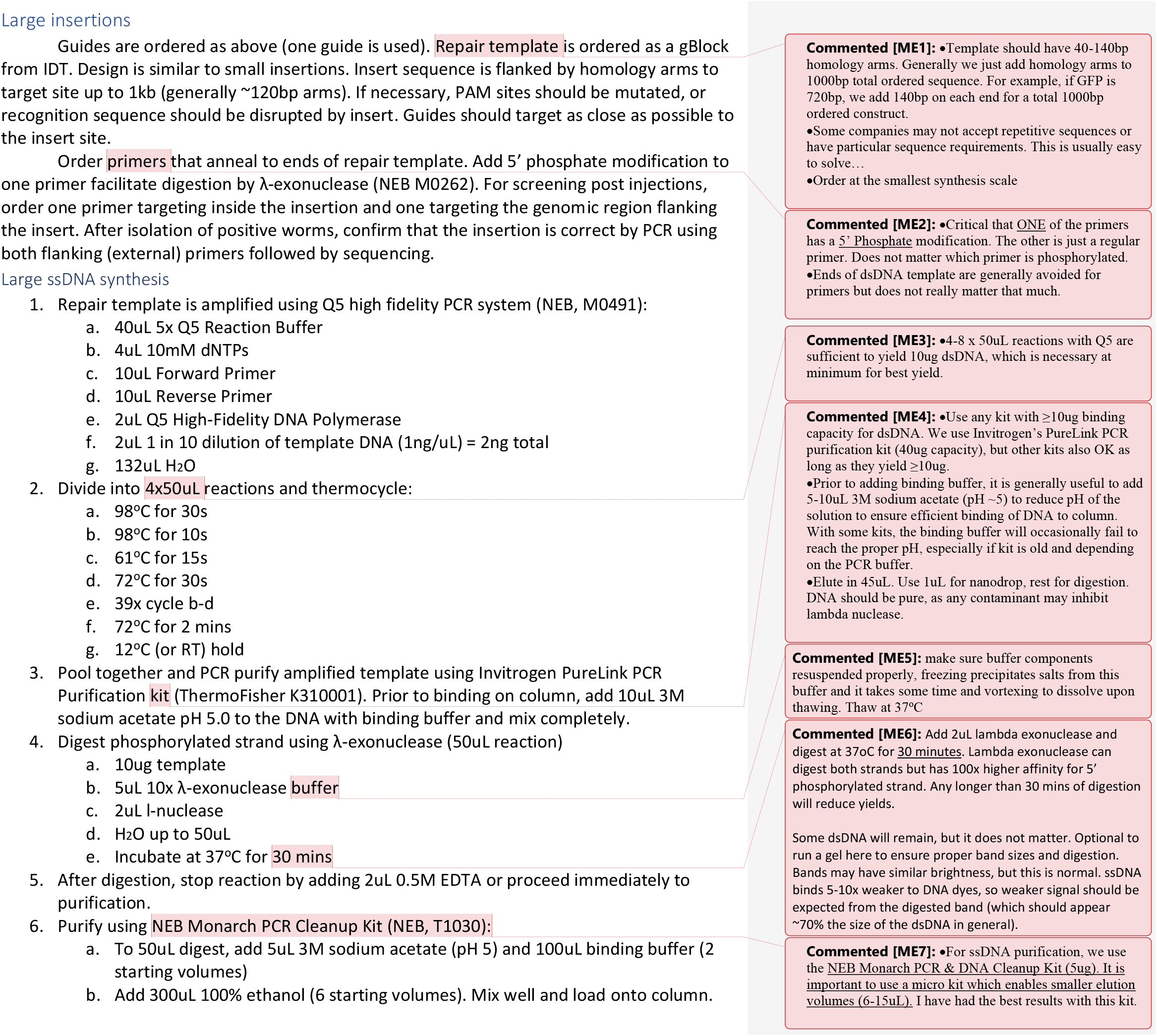
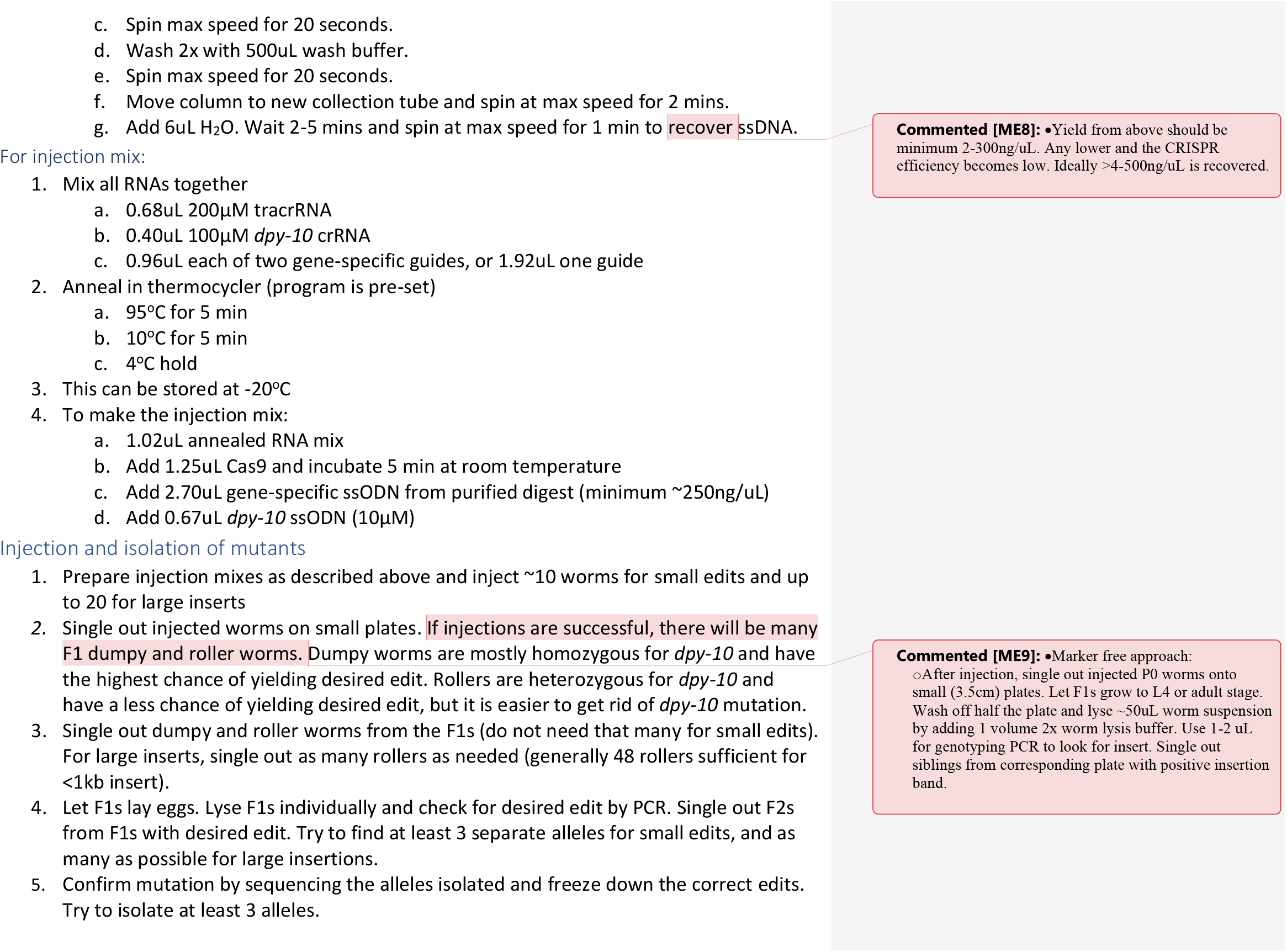

#### Reagents, Commercial Kits, and Oligonucleotides

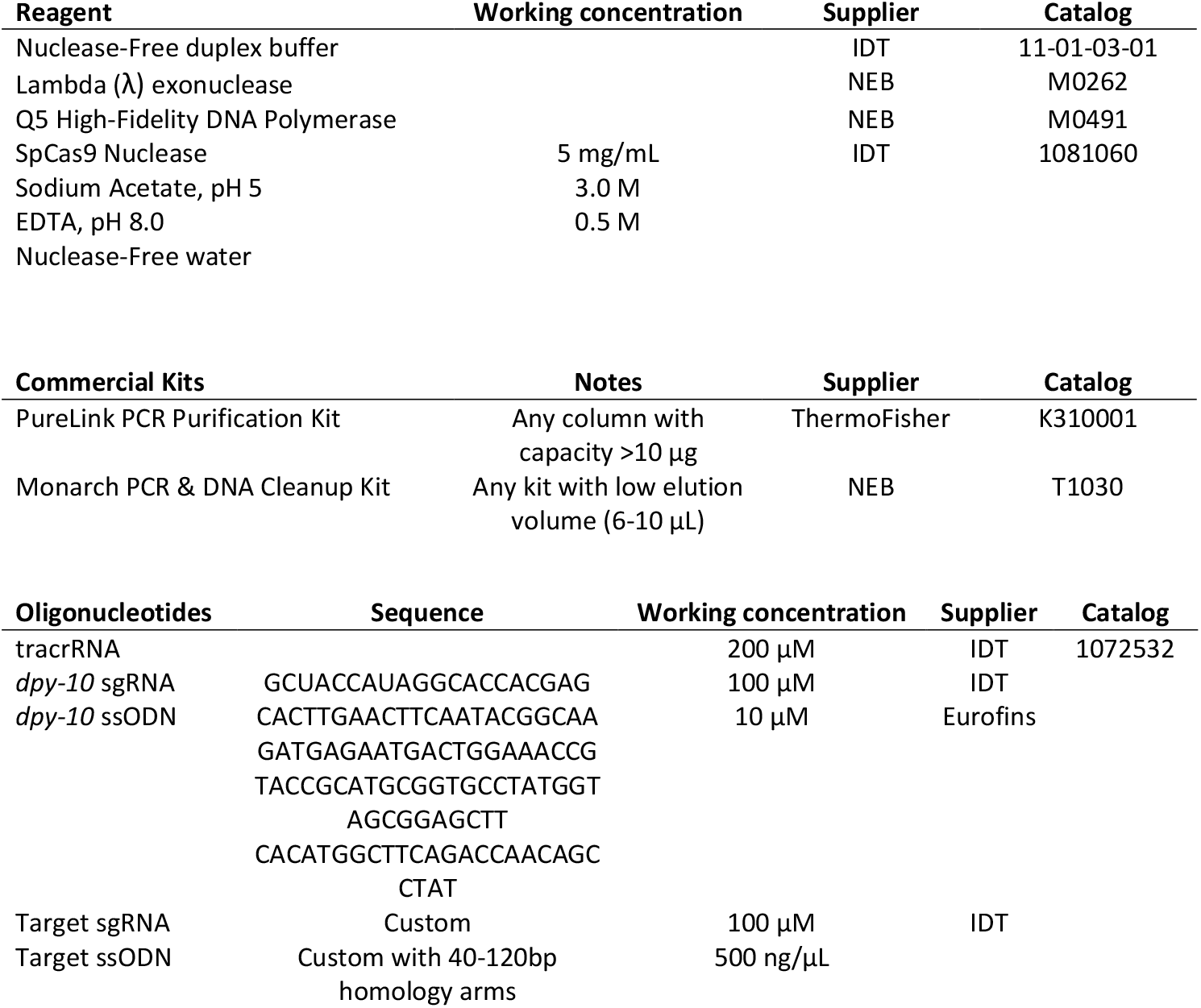

